# The spotlight of attention turns from rhythmic exploration-exploitation to a stable exploitation state

**DOI:** 10.1101/2021.11.18.469122

**Authors:** María Melcón, Sander van Bree, Yolanda Sánchez-Carro, Laura Barreiro-Fernández, Luca D. Kolibius, Elisabet Alzueta, Maria Wimber, Almudena Capilla, Simon Hanslmayr

**Affiliations:** Department of Biological and Health Psychology, Universidad Autónoma de Madrid, Madrid 28049, Spain; Centre for Cognitive Neuroimaging, University of Glasgow, Glasgow G12 8QQ, United Kingdom; Centre for Human Brain Health, University of Birmingham, Birmingham B15 2TT, United Kingdom; Department of Psychiatry, Universidad Autónoma de Madrid, Madrid 28029, Spain; Biosciences Division, Center for Health Sciences, SRI International, Menlo Park, California 94025

**Author notes:** **Corresponding Authors**: María Melcón (first author), Department of Biological and Health Psychology, School of Psychology, Universidad Autónoma de Madrid, C/ Ivan Pavlov 6, 28049, Madrid, Spain, Simon Hanslmayr (senior author), Cognition and Oscillation Lab, Centre for Cognitive Neuroimaging, School of Psychology and Neuroscience, University of Glasgow, 62 Hillhead St, Glasgow G12 8QB, United Kingdom.

**Keywords:** EEG, multivariate pattern analysis, neural decoding, rhythmic cognition, spatial attention

## Abstract

While traditional studies claim that visuospatial attention stays fixed at one location at a time, recent research has rather shown that attention rhythmically fluctuates between different locations at rates of prominent brain rhythms. However, little is known about the temporal dynamics of this fluctuation and, particularly, whether it changes over time. Thus, we addressed this question by investigating how visuospatial attention behaves over space and time. We recorded electroencephalographic activity of twenty-seven human participants while they performed a visuospatial cueing task, where attention was covertly oriented to the left or right visual field. In order to decode the spatial locus of attention from neural activity, we trained and tested a classifier on every timepoint of the orienting period, from the attentional cue to stimulus onset. This resulted in one temporal generalization matrix per participant, which was time-frequency decomposed to identify the sampling rhythm. Finally, a searchlight analysis was conducted to reveal the brain regions responsible for attention allocation. Our results show a dynamic evolution of the attentional spotlight, distinguishing between two states. In an early time window, attention explored both cued and uncued hemifield rhythmically at ~10 Hz. In a later time window attention focused on the cued hemifield. Classification was driven by occipital sources, while frontal regions exclusively became involved just before the spotlight settled onto the cued location. Together, our results define attentional sampling as a quasi-rhythmic dynamic process characterized by an initial rhythmic exploration-exploitation state, which is followed by a stable exploitation state.

## INTRODUCTION

Attention operates as a spotlight, focusing on a single region of space at a time (Eriksen & James, 1986; James, 2007; Posner, 1980). Traditionally, its temporal evolution has been described as constant (Brookshire, 2021; Posner, 1980). In other words, when a stimulus appears, the attentional spotlight (AS) is thought to be deployed to its spatial location, remaining there until it is captured again or voluntarily displaced. However, this idea has been challenged by more recent findings demonstrating that attention is not static but rather rhythmically fluctuates across the visual space (Fiebelkorn et al., 2013a; Landau & Fries, 2012; VanRullen et al., 2007), which has been linked to brain oscillatory dynamics (Buschman & Miller, 2009; Fiebelkorn et al., 2018; Fiebelkorn & Kastner, 2019; Helfrich et al., 2018; Lakatos et al., 2008). These studies distinguish two alternating states in attentional fluctuation: exploitation versus exploration (Fiebelkorn et al., 2018; Gaillard et al., 2020). Exploitation refers to attending to the cued location and it has been associated with top-down signals, whereas exploration consists of attending to the uncued location and is considered a bottom-up process. Overall, evidence comes from recordings of prefrontal areas in non-human primates, although a few works have shown fluctuations also percolate visual regions (Kienitz et al., 2018; Spyropoulos et al., 2018).

What is striking is the wide range of frequencies attentional sampling has been reported, ranging from 3 to 26 Hz, thereby encompassing theta, alpha, and beta rhythms (Buschman & Miller, 2009; Fiebelkorn & Kastner, 2019; Gaillard et al., 2020; Kienitz et al., 2018; Landau & Fries, 2012). One explanation for this variety might be that the rhythm of attentional sampling is not stationary. That is, the frequency at which the AS moves may change over time. Thus, the orienting period could be initially dominated by an exploration-exploitation alternation state that samples the visual field to avoid missing important but unexpected information in a bottom-up manner. Once the visual field has been explored, a stable exploitation state may set in where attentional resources are allocated to the area of interest. This later state would likely be guided by a top-down signal and drastically reduce the frequency of attentional sampling.

This study aims to elucidate the dynamics of the AS using a time-sensitive decoding approach that tracks the location of attention on a millisecond-by-millisecond basis. We use multivariate pattern analysis (MVPA) to decode whether attention at any given timepoint is deployed to the cued or uncued location, which is either the left or the right visual field (LVF or RVF), using high-density electroencephalographic (EEG) recordings in humans.

First, we obtained temporal generalization matrices (TGMs), by training and testing a classifier on every combination of timepoints for both cued hemifields. Such TGMs are ideal to reveal neural code generalizations, that is, neural generators that are reactivated across time (King & Dehaene, 2014). Figure 1 illustrates three hypothetical TGMs according to the scenarios mentioned above, AS as: (i) a constant process (Figure 1A), (ii) a stationary dynamic process (Figure 1B) or (iii) a quasi-rhythmic dynamic process (Figure 1C). The first scenario would give rise to a sustained non-oscillating pattern, in which high accuracy spreads off the diagonal over time (Figure 1A), indicating that one neural pattern tracks the location of attention constantly. The second scenario would lead to rhythmic attentional sampling reversing between locations at a stationary rate. As shown in Figure 1B, the resulting TGM is a yellow-blue checkerboard where the classification criterion at a given training time goes from above chance classification at one testing time to below chance classification at another. However, diagonal performance is above chance; in other words, the AS is decodable online. Under the third scenario, sampling would evolve from shifting between hemifields at a decreasing rate to settling onto the cued hemifield. In terms of the TGM, this would mean a transition from a yellow-blue checkerboard to a large yellow blob (Figure 1C).

**Figure 1.**
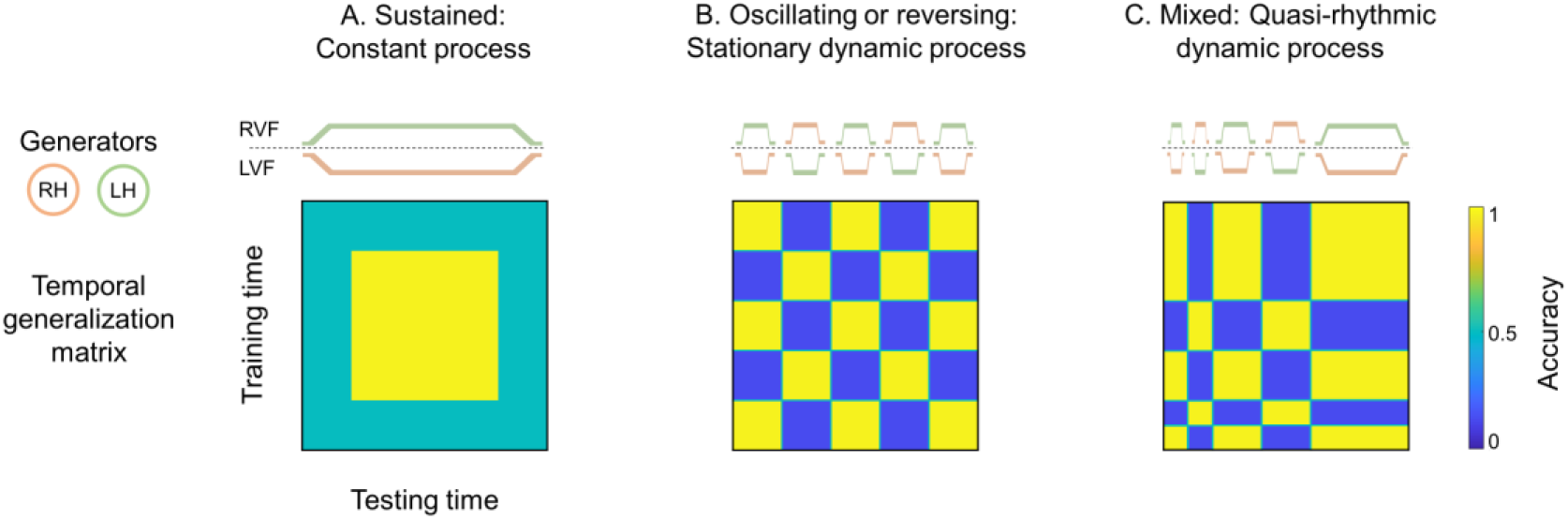
Schematics of different hypothetical scenarios of attentional sampling. The upper panel shows the tracking of the visual field (right, RVF and left, LVF) by the neural generators from the right and left hemispheres (RH and LH) across time. The lower panel depicts the temporal generalization matrix corresponding to each possible scenario. A. Sustained pattern: attentional spotlight (AS) as a constant process, i.e., constant fixation. B. Oscillating pattern: AS as a stationary dynamic process, i.e., sampling information from LVF and RVF at a fixed rhythm. C. Mixed pattern: AS as a quasi-rhythmic dynamic process, i.e., decreasing sampling rate until stabilization.

## MATERIALS AND METHODS

### Participants

Twenty-seven healthy students from the Autónoma University of Madrid with a normal or corrected-to-normal vision took part in this experiment. Data from 3 participants were excluded due to technical problems during EEG recordings, yielding a final sample of 24 participants (20 females; 20 right-handed; mean ± SD age: 19.5 ± 1.9). Students were compensated with course credits for their participation. They all provided written informed consent before the experiment. The Autónoma University of Madrid’s Ethics Committee approved the study, which was conducted in compliance with the Declaration of Helsinki.

### Stimulus material

Stimuli consisted of Gabor patches generated offline using Matlab (R2018a, The MathWorks). Vertically oriented gratings were defined as the target stimuli, with a 10 %probability of appearance, while standard stimuli were angled at ±45°.

Stimuli were presented in 24 locations in the visual field (see Fig. 2A), which was divided according to different polar angles (in 45° steps) and eccentricities (three concentric rings of 2.6°, 9.8° and 22.2° radius), based on Capilla et al. (2016). Stimuli were scaled to the respective 24 locations to compensate for cortical magnification (Baseler et al., 1994; Horton & Hoyt, 1991; Wu et al., 2012).

**Figure 2.**
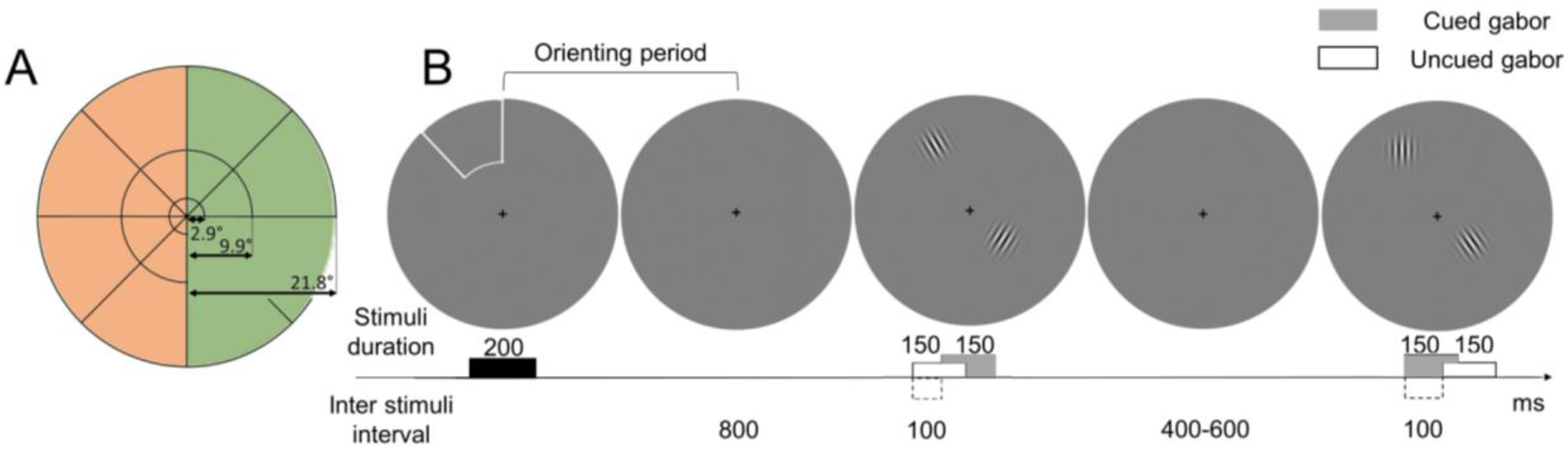
Experimental stimuli and procedure. A. Stimulated locations. The visual field was divided into 24 scaled sectors of different polar angles and eccentricities. Orange and green shades show the experimental conditions: left and right visual field (LVF and RVF, respectively). B. Example of one experimental block. Each block started with an exogenous valid cue. Participants were instructed to covertly pay attention to the cued location and ignore the other locations. After 800 ms, a sequence of 20 pairs of Gabors were presented at both cued and uncued locations. Participants were asked to report cued vertical Gabors. Note that the size of stimuli is not scaled here for visualization purposes.

### Experimental procedure

The study was conducted inside a dimly lit, sound-attenuated and electromagnetically shielded room. The experimental task was carried out using PsychToolbox (v3.0; Brainard, 1997; Kleiner et al., 2007) and presented on a 55-inch monitor, allowing the stimulation of peripheral regions of the visual field. Participants were comfortably seated 80 cm away from the monitor.

Subjects performed a spatially cued discrimination task while maintaining their gaze on a central fixation cross. The task was divided into blocks, the structure of which is depicted in Figure 2B. At the beginning of each block, the border of one sector was illuminated for 200 ms, cueing the location to be attended throughout the block. After an orienting period of 800 ms, a sequence of 20 Gabors pairs was presented for 150 ms each, with a variable inter trial interval ranging from 400 to 600 ms. One of the Gabors in a pair appeared at the cued location while the other was presented at a random uncued location, with a jitter of ±100 ms relative to the previous one. Every sector of the visual field was cued in five blocks, leading to 60 blocks for each hemifield. Stimulation was presented over a gray background.

Participants were instructed to pay covert attention to the cued location while ignoring other locations. They were asked to report target appearance, by pressing a key as quickly and accurately as possible. To avoid fatigue, the experiment was administered in four runs, with self-paced breaks between them. In addition, participants were allowed short breaks between blocks, with the screen turned gray for 3 s. They were encouraged to restrict blinks to these periods, remaining still and relaxed. Before starting the experiment, subjects completed two blocks of practice sessions to familiarize themselves with the task. The experiment lasted approximately 40 minutes.

### EEG recording

The EEG signal was recorded using a BioSemi bioactive electrode cap with 128 electrodes. Online EEG signal was referenced to one external electrode at the nose-tip and sampled at 1024 Hz. We also monitored blinks, vertical and horizontal eye movements by an electrooculogram (EOG) recorded bipolarly from the top and the bottom of the right eye and ~1 cm lateral to the outer canthi of both eyes. Offset of the active electrodes were kept below 25-30 millivolts.

### Data analysis

After acquisition, data preprocessing and analysis was performed in MATLAB using FieldTrip (Oostenveld et al., 2011), MVPA-light (Treder, 2020), and in-house MATLAB code. Overall, the EEG signal analysis aimed to characterize the temporal dynamics of attentional sampling and its neural generators during the orienting period. For this purpose, we first preprocessed the data by removing artifactual activity. Then, we used MVPA to decode the neural footprint of the AS over the time course of the orienting period, i.e., after the cue onset. A classifier trained and tested on every timepoint resulted in one temporal generalization matrix (TGM) for each participant. To identify the rhythm of the attentional sampling, we performed time-frequency analyses on these matrices. Finally, to localize which brain regions contributed most to the classification, we used a searchlight analysis on the scalp and the source level. This procedure is described in more detail in the following sections.

#### Preprocessing

The continuous EEG signal was segmented into 4500 ms epochs, starting 2000 ms prior to cue onset. The use of these long epochs allowed us to avoid edge artifacts in later time-frequency analyses. Note that only periods immediately after the cue onset were analysed since the study of the brain response induced by target and distractor stimuli is beyond the aim of this work. Data were low-pass filtered below 30 Hz, baseline corrected (−200 to 0 ms) and re-referenced to the common average. We did not high-pass filter the data to avoid decoding artefacts in the TGM (van Driel et al., 2021). Finally, time delays between expected and actual presentation of visual stimuli on the screen were corrected employing a photodiode.

Artifact rejection was carried out in a three-step procedure. First, we eliminated trials with eye movements by visual inspection, as they may indicate a failure to maintain covert attention. Secondly, we used Independent Component Analysis (ICA) to remove remaining artifacts, related to eye-blinks or muscular activity. Finally, bad channels were interpolated using data of adjacent electrodes (mean ± SD of interpolated electrodes: 1.2 ± 2.7).

After artifact rejection, data were down-sampled to 128 Hz to reduce computational time and split into two experimental conditions based on the cued location: LVF or RVF.

#### Temporal generalization analysis

To decode the neural behaviour of attention in the visual field, we used the temporal generalization method (King & Dehaene, 2014). Before classification, we kept trial numbers between conditions constant by random selection, which resulted in an average of 113.7 ± 13.7 (mean ± SD) epochs per participant. Then, a linear discriminant analysis (LDA, (Fisher, 1936) was used as a classifier. This classifier was trained and tested for each participant on every timepoint by using the z-scored amplitude from the 128 EEG channels across trials as features. This resulted in a temporal generalization matrix (TGM) with one value for every combination of training and testing times. A TGM not only indicates when brain activity differs between experimental conditions, but also if a neural pattern present at a given timepoint is reactivated at any other time, shedding light on the temporal organization of information processing. To avoid overfitting, this procedure was applied following a cross-validation scheme (Lemm et al., 2011) where the data were randomly split into five folds. Four of these folds were first used as the training set to find the decision boundary that best separated both classes. Then, that boundary was applied to the fifth fold, i.e., the testing set, to predict which class the neural pattern belonged to. The process was repeated until each fold was used as the testing set once. In addition, to avoid obtaining a biased result due to the random folding and to ensure a stable frequency measurement in later steps, the analysis was repeated 15 times with new random folds and then averaged across those 15 runs.

The classification metric was accuracy, which is the fraction of trials correctly predicted by the classifier (being 50% considered chance level). We used a non-parametric two-level permutation approach to test when the classifier performance was statistically significant. To this end, we first created 50 permuted TGMs per participant by randomly shuffling the classification labels (LVF and RVF). We z-scored both real and shuffled TGMs for each participant according to the mean and standard deviation of their shuffled TGMs and used these z-scored TGMs in all subsequent steps as well as in the further frequency analysis. Then, we generated a random distribution for each testing time by training time combination over 10,000 iterations. In each repetition, we randomly drew one of the 50 z-scored shuffled TGMs per participant and computed a grand-average TGM (i.e., mean of the 24 randomly selected TGMs) under the null hypothesis. The resulting distribution of the means was used to derive the empirical p-values of the grand-average real TGM. Thus, p-values below/above the 2.5^th^/97.5^th^ percentiles in each point of the TGM were considered statistically significant. P-values were previously corrected for multiple comparisons by False Discovery Rate (Benjamini & Hochberg, 1995).

#### Time-frequency analysis

We then conducted a time-frequency analysis to identify the rate at which attentional sampling occurs throughout time. For this purpose, each participant’s TGM was decomposed using a Hanning-tapered sliding window Fourier transform. Oscillatory power was estimated from 0 to 1 s for each time sample (steps of 7.8 ms) at 1 Hz steps between 2 and 30 Hz. The width of the Hanning window was adjusted to 5 cycles per frequency. Note that we had segmented the data into 4.5 s epochs to avoid edge artifacts and that we had z-scored the individuals TGMs according to each participant shuffled mean and standard deviation.

This analysis was applied independently to each temporal series of the TGM (i.e., to each row and column; see Figure 3, left), leading to two three-dimensional matrices: training time x testing time x frequency. On the one hand, the rows matrix stored the power decomposition of each frequency in the same training time and along testing time (Figure 3, top-middle). On the other hand, the columns matrix stored the opposite result, i.e., power decomposition in the same testing time along the training time (Figure 3, bottom-middle). Subsequently, both matrices were averaged resulting in a single three-dimensional power matrix of dimensions training time x testing time x frequency (Figure 3, right). In this way, we obtained a single value for each voxel where the power of the rows and columns was equally represented.

**Figure 3.**
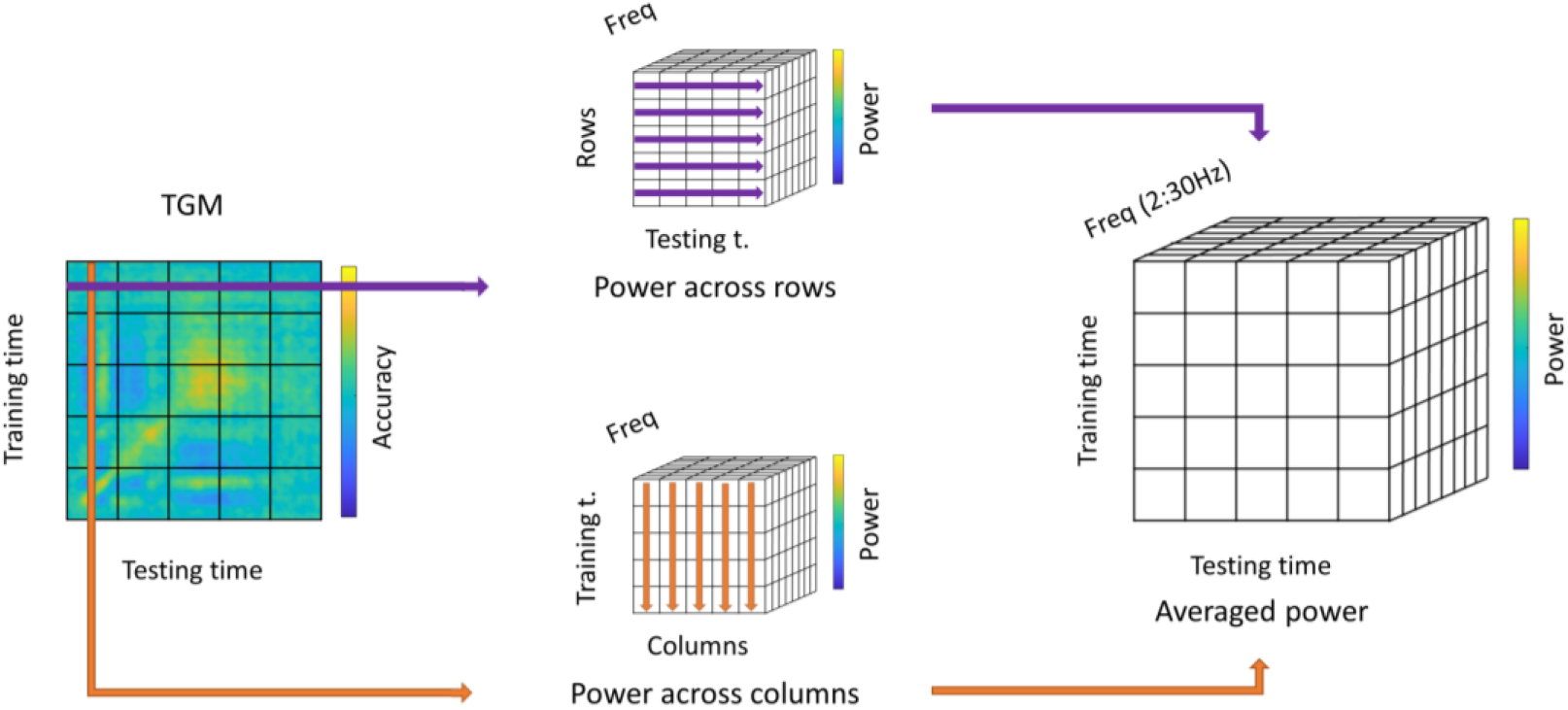
Time-frequency analysis of the TGM. We applied a time-frequency analysis independently over each row and column of the TGM (left). This resulted in two power matrices, one for the testing time evolution (top-middle) and another one for the training time evolution (bottom-middle). Both matrices were averaged to yield one composite matrix that quantifies the distribution of classifier frequencies in the TGM (right).

Finally, to statistically test the results of the time-frequency analysis, we used the same non-parametric two-level permutation approach as in the previous section. Time-frequency representations of the 50 z-scored permuted TGMs for each participant were computed as explained above. Over 10,000 iterations, one shuffled power TGM per participant was randomly selected in each repetition to generate a grand-average TGM. This led to an empirical distribution where FDR corrected p-values below/above the 2.5/97.5^th^ percentiles in each point and frequency of the TGM were considered statistically significant. We selected the significant dominant frequency peak at each timepoint, reducing the matrix to two dimensions. Then, this output was combined with the statistical result of the classifier performance, so that the resulting masked TGM in Figure 4C shows the significant dominant frequency peaks at timepoints where classifier performance was above chance level.

**Figure 4.**
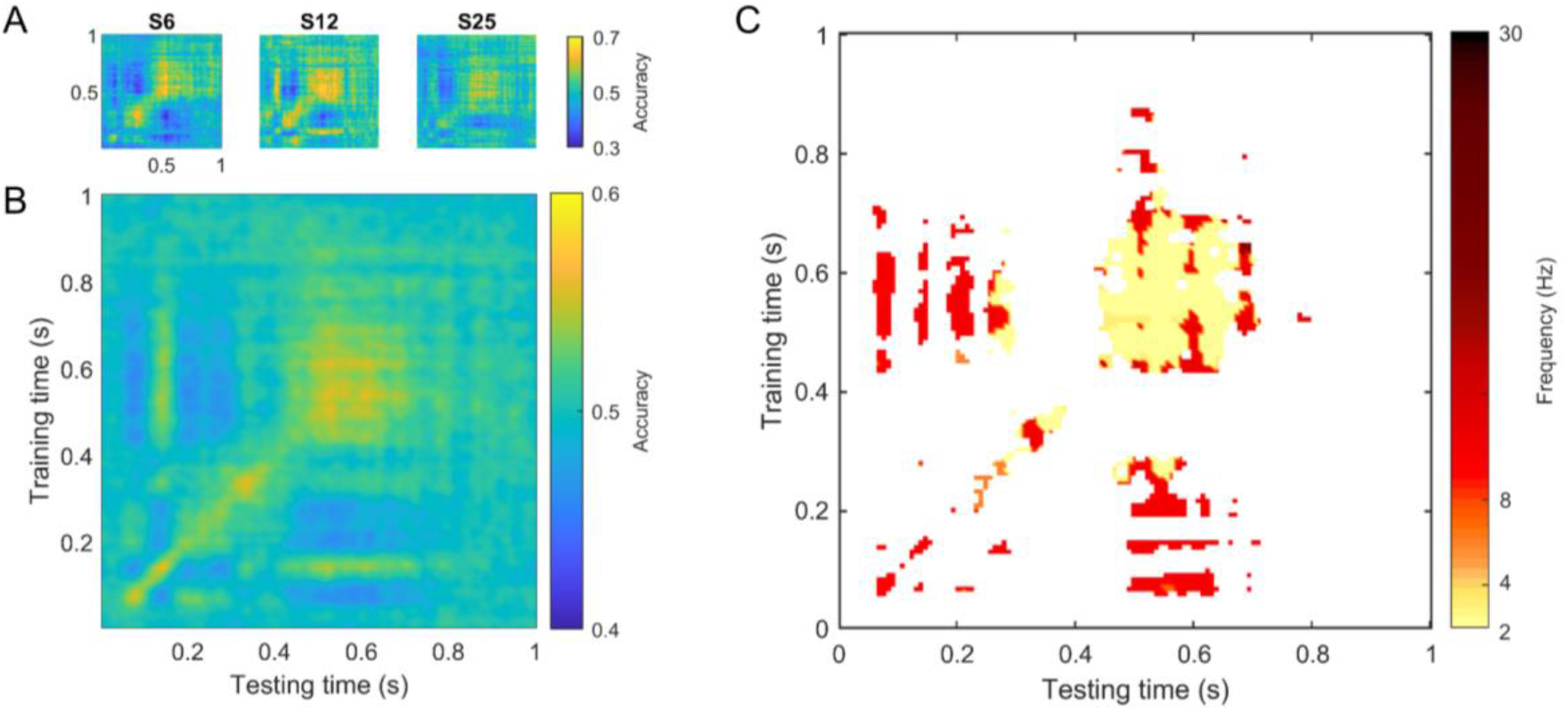
Temporal classification results. A. Temporal generalization matrix for 3 individual participants. B. Temporal generalization matrix averaged across all participants. C. Significant dominant sampling rhythm thresholded by significant classification accuracy (both FDR corrected).

#### Searchlight analysis

Statistical analysis of the classifier performance revealed four significant time windows across the TGM diagonal (around 65-95, 110-165, 315-380 and 480-660 ms). In order to identify the brain regions that contained critical information to discern between LVF and RVF in these time windows, we conducted a searchlight analysis (Kriegeskorte et al., 2006). In this approach, a classifier is trained on a subselection of the data (i.e., centered on neighbouring sensors or voxels) which is termed the searchlight. This searchlight is then systematically shifted across the whole brain/sensor space to yield one decoding performance value per region. To this end, we first set the cross-validation parameters to 5 folds and 15 repetitions and defined the size of the searchlight, considering each electrode and its three nearest neighbouring channels. Then, classification was performed independently for each significant time window (65-95, 110-165, 315-380 and 480-660 ms), leading to four accuracy maps across the scalp.

Finally, we applied the same procedure in source space (see below) to identify the underlying brain sources. We selected the voxels inside the brain as features (1165 voxels) and conducted the search by moving a 3-cm radius sphere throughout that brain volume. This resulted in four analogue 3D brain maps of classification accuracies.

#### Source analysis

The localization of brain sources was performed by means of beamforming (Gross et al., 2001; Van Veen et al., 1997). We estimated the activity generated by each voxel of a standard magnetic resonance image (MRI) in MNI (Montreal Neurological Institute) space by employing a standard boundary element method (BEM) volume conduction model (Oostenveld et al., 2003) and standard 10-05 electrode positions. After segmenting the MRI into 12-mm voxels, we computed the lead fields corresponding to the 3 possible orientations for each of them. We used our time window of interest (from 0 to 1 s) to calculate the single-trial covariance matrix, which was then used to compute the spatial filter coefficients by means of linearly constrained minimum variance beamformer (LCMV, Van Veen et al., 1997). The regularization parameter lambda was set to 10%.

Subsequently, we projected each sensor-level trial into each voxel of source-space employing the spatial filter corresponding to the optimally oriented dipole. To remove the centre of the head bias, we normalized source-level activity at each voxel by dividing by the root-mean-square across trials.

#### Code accessibility

The data and scripts to reproduce the analysis and figures in this paper are available on the Open Science Framework platform (https://osf.io/8e9c7/).

## RESULTS

### Behavioural results

Hit rates for trials in which participants were covertly oriented to the LVF (36.5 ± 10%, mean ± SD) or RVF (37.1 ± 10.9%, mean ± SD) did not statistically differ (t(23) = −0.41, p = 0.68). Likewise, reaction times for correct trials were not different (t(23) = −0.00, p = 0.99) for LVF (547 ± 52 ms, mean ± SD) and RVF (547 ± 56 ms, mean ± SD).

### Temporal evolution of the attentional spotlight

We employed a multivariate temporal classification approach to examine whether the AS samples information between LVF and RVF. For this purpose, we trained and tested a classifier to distinguish the two cued hemifields on every combination of timepoints. As illustrated in Figure 1, a constant spotlight would be reflected in a sustained pattern, a stationary rhythmic spotlight would lead to an oscillating pattern, and a quasi-stationary dynamic spotlight would lead to a combination of the above, with the frequency of the attentional fluctuation decreasing over time.

Representative individual TGMs are shown in Figure 4A which revealed that the AS behaves in a fluctuating dynamic way. This pattern was also evident in the grand-average TGM (Figure 4B) which clearly shows that classifier accuracy fluctuated between 0.4 and 0.6 (Figure 4B). Furthermore, the non-parametric permutation test demonstrated that classification accuracy differed significantly from the chance level at various points in the TGM (see Figure 4C). Importantly, significant results were obtained for both accuracies below and above chance level (see blue and yellow areas in Fig. 4B, respectively). Below chance level accuracies were confined to the off-diagonal areas of the TGM, with the diagonal showing only above chance level classification accuracies. The fluctuation of the AS is particularly clear in the early time window (between 50 and 400 ms), and less present at later timepoints (after ~700 ms).

Interestingly, the left and bottom borders of the TGM show a vertical and horizontal structure respectively, with a stable pattern being present approximately from 400 to 700 ms on both time axes. This indicates that the early transient attentional state generalized to a later, apparently more sustained state, suggesting that the neural activity pattern that is present around 150 ms reoccurs in the late, sustained period. Thus, attention first appears to sample information rhythmically from both hemifields, before it settles steadily onto the cued location. These results suggest that the AS would have two sequential modes of operation, moving back and forth between hemifields during an early exploration-exploitation stage before settling towards the cued hemifield during a later single exploitation phase. Therefore, these data support the third of the three scenarios outlined in Figure 1.

To investigate the temporal evolution of attentional sampling more systematically, we obtained the time-frequency decomposition of the classifier performance. We applied this analysis to each row and column of the TGM and determined the significant peak frequency when the classifier remained above chance level. If attention sampled information rhythmically but at a decreasing rate over time, then the dominant frequency of the TGM should drop across time. As shown in Figure 4C, this is exactly what can be observed. During the first 200 ms in both training and testing time, attention fluctuated at a rate of 10 Hz. As time goes on, the fluctuation rate decreased, down to 2 Hz at around 500 ms. Although these were the most dominant frequencies, it is noteworthy that almost all of the frequencies of attentional sampling mentioned in previous studies can be observed here at different time points (i.e., from 3 Hz up to 26 Hz, (Buschman & Miller, 2009; Fiebelkorn & Kastner, 2019; Gaillard et al., 2020; Kienitz et al., 2018; Landau & Fries, 2012).

### Brain regions underlying attentional sampling

A searchlight analysis (Kriegeskorte et al., 2006) was conducted to reveal which brain regions contributed most to the decoding of the actual location of the AS. For this purpose, we selected four time windows as revealed by the statistical analysis of the TGM (around 65-95, 110-165, 315-380 and 480-660 ms, where classification was above chance level on the diagonal). For each participant and time window, we applied a moving mask of a sphere of three neighbouring channels for the sensor-level analysis, and a sphere of 3-cm radius for source-level. Then, we fitted a linear model independently for each sphere to predict left and right cue activity.

As shown in Figure 5, sensor and source-space analysis led to similar results. Overall, the main generators of significant classification accuracy were localised over visual and frontal regions. Visual areas contributed to classifier performance at early and later time windows, probably indexing the actual locus of the AS. These visual regions spread medially across both hemispheres, showing the peak voxel in source-space over the left hemisphere. In contrast, the contribution of frontal regions, particularly the right ventral frontal area, was confined to an intermediate time window (315-380 ms). Interestingly, activity on prefrontal areas was observed right before the AS settled onto the cued location, which might be indicative of a top-down signal that stops sampling between hemifields, giving rise to the exploitation stage.

**Figure 5.**
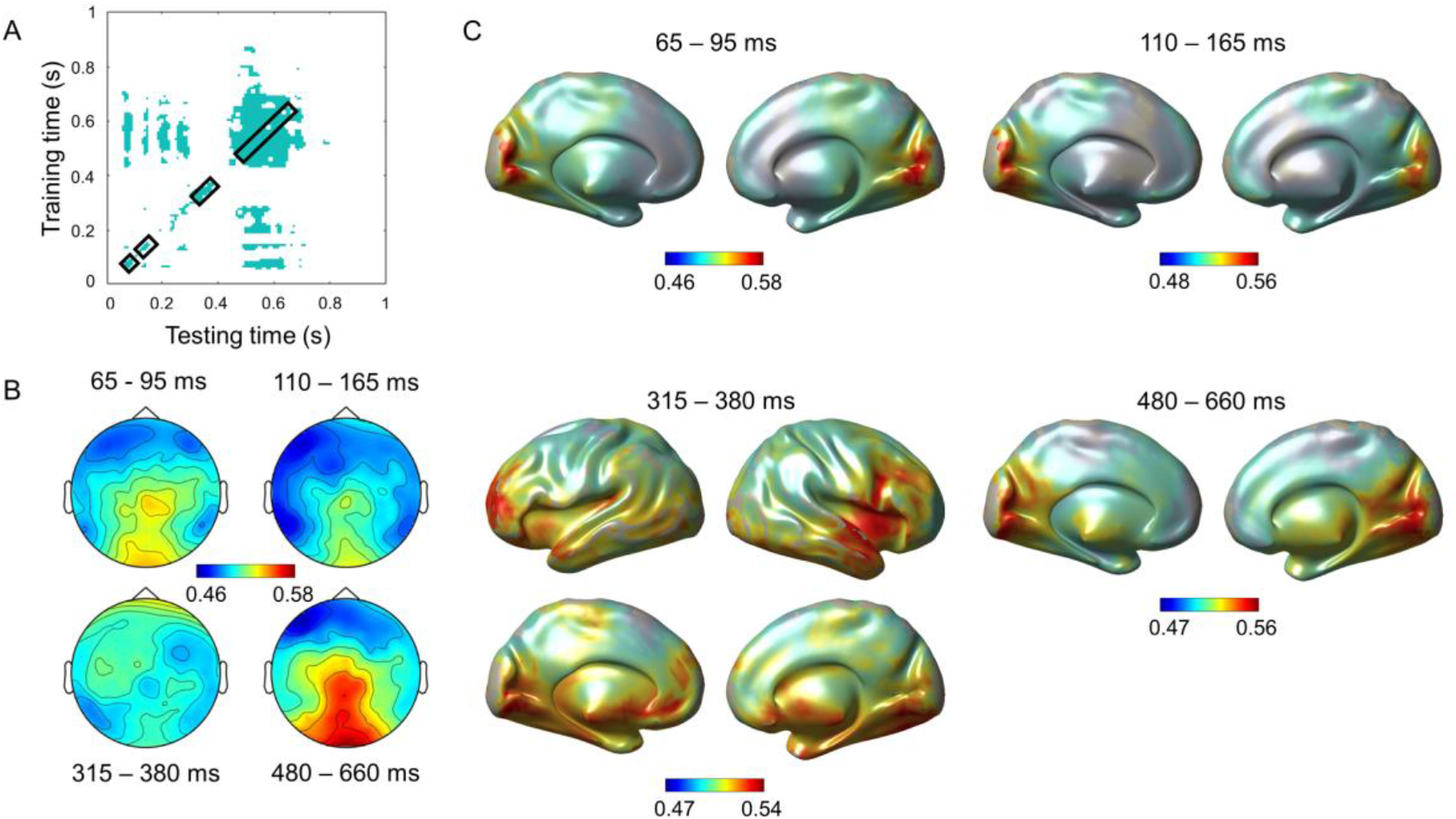
Searchlight results. A. Significant mask for the classifier performance. Squares along the diagonal indicate the time windows on which searchlight analyses were applied. B. Searchlight results in sensor-space for each time window. C. Searchlight results in source-space.

## DISCUSSION

This study aimed to shed light on the spatiotemporal dynamics of the AS. Novel frequency analysis of classifier performance revealed a non-stationary quasi-rhythmic process. This process started by sampling information rhythmically between cued and uncued locations at a rate of ~10 Hz before settling onto the cued location. The earlier process might reflect a bottom-up exploration-exploitation process, whereas the latter process might index a top-down driven exploitation state. Searchlight analysis pointed to visual areas as critical to track the actual locus of the AS in early as well as later stages, while frontal regions might be rather involved in the switch from early alternation to later exploitation.

Previous literature has defined the AS as either a constant process in space and time (Posner, 1980) or a stationary oscillatory process (Fiebelkorn et al., 2013; Landau & Fries, 2012; VanRullen et al., 2007), which would sample information alternately from both hemifields at a wide range of frequencies (Buschman & Miller, 2009; Fiebelkorn & Kastner, 2019; Gaillard et al., 2020; Kienitz et al., 2018; Landau & Fries, 2012). Thus, recent studies assume the existence of two states in the sampling behaviour of the AS: exploitation/sampling versus exploration/shifting (Fiebelkorn et al., 2018; Gaillard et al., 2020). Exploitation refers to a phase where attention is being located to the cued position triggered by a top-down signal, while exploration refers to sampling information from uncued locations and reflects a bottom-up driven process. Our results show that both processes are at work in an attentional cueing paradigm but at different times, with an initial rhythmic fluctuation of attention at ~10 Hz, which subsequently, slows down to 2 Hz. This slowing of the sampling frequency is a central observation as it characterizes how the brain strikes a balance between the processes of exploration-exploitation and single exploitation. More specifically, the exploration-exploitation state takes place during the first 400 ms after the cue, where attention moves quickly back and forth between hemifields at a rate of ~10 Hz. This would ensure that no important but unexpected information is being missed. Then a more stable phase settles in where attention remains focused on the cued hemifield, from 400 to 800 ms. This would be consistent with a state of exploitation where the information presented by the cue is being maximally extracted.

The combination of a highly time-sensitive whole-brain recording method (i.e., 128-channel EEG) with a powerful MVPA algorithm enabled us to identify the spatiotemporal dynamics of the AS. For the current findings to emerge it was critical to consider the TGM as a whole, instead of applying the common approach of only investigating accuracy on the diagonal. Indeed, if we had only considered the diagonal, we would have entirely missed the alternating sampling behaviour of the AS (i.e., rhythmically going back and forth between the two hemifields). Our results show that the below-chance classification accuracy on the off-diagonals are crucial since it indicates a rhythmic reversal of hemifield states during the ~400 ms after cue onset. This in turn suggests the existence of a rhythmic attentional sampling at about 10 Hz, similar to the sampling frequencies reported in some previous studies (Dugué et al., 2016; Fiebelkorn et al., 2013b; Landau et al., 2015; Voloh et al., 2015). However, other studies have reported disparate sampling frequencies ranging from 3 to 26 Hz (Buschman & Miller, 2009; Fiebelkorn & Kastner, 2019; Gaillard et al., 2020; Kienitz et al., 2018; Landau & Fries, 2012). Noteworthy, most of these frequencies were observed in the TGM computed in the present study. Our results may help clarifying such inconsistency, by demonstrating that attentional sampling is not a stationary phenomenon but instead changes across time.

An additional remark concerns cue validity. Previous studies describing the AS as a stable oscillatory process have used both informative and non-informative cues (Fiebelkorn et al., 2018; Gaillard et al., 2020; Landau & Fries, 2012). By using time-sensitive decoding methods, the current study shows that the AS is a quasi-rhythmic dynamic process when a 100% valid informative cue is employed. However, the expectations generated by cue validity (Carrasco et al., 2002), might lead to different attentional behaviour. Thus, the single exploitation state might have been triggered by the high predictive value of the cue, whereas attentional sampling may remain constant for intermediate values of cue validity. Nevertheless, we cannot rule out the opposite possibility, that attentional sampling is a quasi-rhythmic dynamic process irrespective of cue validity. Thus, the attentional system might initially behave as a sampling process that dies out over time, ultimately leading to the settling of attention onto the cued location regardless of the information provided by the cue. This possible dichotomy on the behaviour of the AS depending on cue validity is interesting and worthy of further investigation.

Previous studies have found that visual areas, frontal eye-fields and lateral intraparietal sulcus are involved in attention allocation (Fiebelkorn et al., 2018; Gaillard et al., 2020; Kienitz et al., 2018; Spyropoulos et al., 2018). All these works have been carried out in a hypothesis-driven manner, making local recordings from selected brain areas of interest in non-human primates. In contrast, we recorded whole-brain measurements by high-density EEG in human volunteers and employed a data-driven approach (i.e., MVPA) to analyse brain activity. Interestingly, the brain regions underlying attention allocation revealed by our study are similar to those reported in the literature. An occipital source appeared as the area with the largest contribution during the whole epoch. That is, visual cortex was not only recruited during the exploration-exploitation alternating period, but it was also involved in the single exploitation state. This suggests that visual areas track the actual locus of attention, pointing towards the existence of an occipital AS. In addition, it is worth mentioning that classification mainly relied on information from left occipital regions, which fits with the well-known right hemisphere dominance for attention (Heilman & Van Den Abell, 1980). If the right hemisphere is involved in attending to both hemifields but the left hemisphere is exclusively engaged by the contralateral hemifield, only the latter will be informative for the classifier to distinguish between the experimental conditions as we observed here. An interesting and unexpected finding was the activation (i.e., by means of driving classifier performance) of right ventral frontal areas after the exploration stage. We can speculate that a higher-order signal is needed for the occipital AS to stop sampling from both hemifields in order to settle onto the cued location. This would be consistent with the idea that the prefrontal cortex is at the origin of an attentional control signal (Buschman & Miller, 2007; Gregoriou et al., 2014; Moore & Fallah, 2004), and in particular with studies implicating the right ventral prefrontal cortex in inhibitory control (Aron et al., 2003; Chambers et al., 2007). However, our results point that this domain-general control region is multivariately but not univariately involved, as it makes it possible to distinguish between experimental conditions. This could be explained by the ability of the frontoparietal network to adaptively represent task-relevant information (Duncan, 2001; Duncan, 2010). Specifically, the firing pattern of single neurons in this network is adjusted to the specific information relevant for the task, including the location that triggers attentional sampling, as occurs in our results. Indeed, a relationship between the frontoparietal network and the coding of visual information has already been shown (Woolgar et al., 2016). Finally, our analysis revealed no involvement of the parietal lobe, a key region in spatial attention (Critchley, 1953). Its absence makes sense, however, since the parietal lobe doesn’t carry information about the locus of attention, but it is involved in attentional orientation deployment (Posner & Dehaene, 1994). In other words, it shows similar activity across both classes (i.e. left vs right visual field) and therefore would not be picked up by the classifier as employed here.

Together, our results go one critical step beyond previous attempts to characterise the spatiotemporal dynamics of visuospatial attention, revealing a quasi-rhythmic dynamic process. After cue onset, an automatic rhythmic shifting response at ~10 Hz is triggered. Thus, attention would initially alternate between both hemifields, being the allocation of the AS tracked by visual areas. This default mode of the attentional system might be interpreted as a preparation of the visual areas to avoid missing out any potentially important information in the visual field, striking a balance between exploration and exploitation. However, for attention to be effective, it needs to be under top-down control (i.e., it needs to be controlled voluntarily to sample information from areas of interest). Hence, once cue information has been processed, right frontal areas may fixate sampling in the cued hemifield, leading to a later single exploitation stage tracked again by visual areas. However, it should be noted that our data analysis approach focuses directly on neural representations, but not on the brain oscillatory activity underlying these processes. Therefore, the oscillatory mechanisms underlying attentional orientation will have to be disentangled in future studies.

## Acknowledgments

This work was supported by FEDER/Ministerio de Ciencia, Innovación y Universidades – Agencia Estatal de Investigación, Spain (grant PGC2018-100682-B-I00) and the Research Grant FPI-UAM 2017 (UAM, Spain). We are thankful to Ian Charest for valuable advice regarding the use of linear discriminant analysis.

